# The mechanism of transcription start site selection

**DOI:** 10.1101/156877

**Authors:** Libing Yu, Jared T. Winkelman, Chirangini Pukhrambam, Terence R. Strick, Bryce E. Nickels, Richard H. Ebright

## Abstract

During transcription initiation, RNA polymerase (RNAP) binds to promoter DNA, unwinds promoter DNA to form an RNAP-promoter open complex (RPo) containing a single-stranded "transcription bubble," and selects a transcription start site (TSS). TSS selection occurs at different positions within the promoter region, depending on promoter sequence and initiating-substrate concentration. Variability in TSS selection has been proposed to involve DNA "scrunching" and "anti-scrunching," the hallmarks of which are: (i) forward and reverse movement of the RNAP leading edge, but not trailing edge, relative to DNA, and (ii) expansion and contraction of the transcription bubble. Here, using *in vitro* and *in vivo* protein-DNA photocrosslinking and single-molecule nanomanipulation, we show bacterial TSS selection exhibits both hallmarks of scrunching and anti-scrunching, and we define energetics of scrunching and anti-scrunching. The results establish the mechanism of TSS selection by bacterial RNAP and suggest a general mechanism for TSS selection by bacterial, archaeal, and eukaryotic RNAP.

## INTRODUCTION

During transcription initiation, RNA polymerase (RNAP) holoenzyme binds to promoter DNA making sequence-specific interactions with core promoter elements, unwinds a turn of promoter DNA to form an RNAP-promoter open complex (RPo) containing an unwound "transcription bubble," and selects a transcription start site (TSS). The distance between core promoter elements and the TSS can vary, depending upon promoter sequences and concentrations of initiating substrates (Jeong and Kang, 1994; Liu and Turnbough, 1994; Walker and Osuna, 2002; Lewis and Adhya, 2004; Vvedenskaya et al., 2015; Winkelman et al., 2016a,b). We previously have proposed that variability in the distance between core promoter elements and the TSS is accommodated by DNA "scrunching" and "anti-scrunching," the defining hallmarks of which are: (i) forward and reverse movements of the RNAP leading edge, but not the RNAP trailing edge, relative to DNA and (ii) expansion and contraction of the transcription bubble (Vvedenskaya et al., 2015; Winkelman et al., 2016a,b).

## RESULTS AND DISCUSSION

### TSS selection exhibits first hallmark of scrunching--movements of RNAP leading edge but not RNAP trailing edge--both *in vitro* and *in vivo*

In prior work, we demonstrated that bacterial TSS selection *in vitro* exhibits the first hallmark of scrunching by defining, simultaneously, the TSS, the RNAP leading-edge position, and RNAP trailing-edge position for transcription complexes formed on a library of 10^6^ promoter sequences (Winkelman et al., 2016a). We used RNA-seq to define the TSS, and we used unnatural-amino-acid-mutagenesis, incorporating the photoactivatable amino acid *p*-benzoyl-L-phenylalanine (Bpa), and protein-DNA photocrosslinking to define RNAP-leading-edge and trailing-edge positions (Winkelman et al., 2016a). The results showed that the “discriminator” element, the 3 nt sequence located immediately downstream of the promoter −10 element (Haugen et al., 2006; Feklistov et al. 2006), is a determinant of TSS selection. The results further showed that, as the TSS changes for different discriminator sequences, the RNAP-leading-edge position changes, but the RNAP-trailing-edge position does not change. For example, replacing a GGG discriminator by a CCT discriminator causes a 2 bp downstream change in TSS (from the position 7 bp downstream of the −10 element to the position 9 bp downstream of the −10 element), causes a 2 bp downstream change in RNAP leading-edge position, but does not cause a change in RNAP trailing-edge position (Fig. 1A).

To determine whether bacterial TSS selection *in vivo* also exhibits the first hallmark of scrunching, we adapted the above unnatural-amino-acid-mutagenesis and protein-DNA-photocrosslinking procedures to define RNAP leading-edge and trailing-edge positions in TSS selection in living cells (Figs. 1, S1–S2). We developed approaches to assemble, trap, and UV-irradiate RPo formed by a Bpa-labeled RNAP derivative in living cells, to extract crosslinked material from cells, and and to map crosslinks at single-nucleotide resolution (Fig. S1). In order to assemble, trap, and UV-irradiate RPo in living cells, despite the presence of high concentrations of initiating substrates that rapidly convert RPo into transcribing complexes, we used a mutationally inactivated RNAP derivative, β’D460A, that lacks a residue required for binding of the RNAP-active-center catalytic metal ion and initiating substrates (Zaychikov et al., 1996) (Figs. S1–S2). Control experiments confirm that, *in vitro,* in both the absence and presence of initiating substrates, the mutationally inactivated RNAP derivative remains trapped in RPo, exhibiting the same pattern of leading-edge and trailing-edge crosslinks as wild-type RNAP in the absence of initiating substrates (Fig. S2). In order to introduce Bpa at the leading-edge and trailing-edge of RPo in living cells, we co-produced, in *Escherichia coli,* a Bpa-labeled, decahistidine-tagged, mutationally inactivated RNAP derivative in the presence of unlabeled, untagged, wild-type RNAP, using a three-plasmid system comprising (1) a plasmid carrying a gene for RNAP β’ subunit that contained a nonsense codon at the site for incorporation of Bpa, the β’D460A mutation, and a decahistidine coding sequence; (2) a plasmid carrying genes for n engineered Bpa-specific nonsense-suppressor tRNA and an engineered Bpa-specific aminoacyl-tRNA synthase (Chin et al., 2002); and (3) a plasmid containing a promoter of interest (Fig. S1A). (Using this merodiploid system, with both a plasmid-borne mutant gene for β’ subunit and a chromosomal wild-type gene for β’ subunit, enabled analysis of the mutationally inactivated RNAP derivative without loss of viability.) In order to perform RNAP-DNA crosslinking and to map resulting RNAP-DNA crosslinks, we grew cells in medium containing Bpa, UV-irradiated cells, lysed cells, purified crosslinked material using immobilized metal-ion-affinity chromatography targeting the decahistidine tag on the Bpa-labeled, decahistidine-tagged, mutationally inactivated RNAP derivative, and mapped crosslinks using primer extension (Fig. S1B). The results showed an exact correspondence of crosslinking patterns *in vitro* and *in vivo* (Fig. 1B, “*in vitro*” vs. “*in vivo*” lanes). The RNAP leading edge crosslinked 2 bp further downstream on CCT than on GGG, whereas the RNAP trailing edge crosslinked at the same positions on CCT and GGG (Fig. 1B). We conclude that TSS selection *in vivo* shows the first hallmark of scrunching.

### TSS selection exhibits second hallmark of scrunching--changes in size of transcription bubble

The results in Fig. 1 establish that TSS selection *in vitro* and *in vivo* exhibits the first hallmark of scrunching. However, definitive demonstration that TSS selection involves scrunching also requires demonstration of the second hallmark of scrunching: i.e., changes in transcription-bubble size. To determine whether bacterial TSS selection exhibits the second hallmark of scrunching we used a magnetic-tweezers single-molecule DNA-nanomanipulation assay that enables detection of RNAP-dependent DNA unwinding with near-single-base-pair spatial resolution and sub-second temporal resolution (Revyakin et al., 2004, 2005, 2006) to assess whether TSS selection correlates with transcription-bubble size for GGG and CCT promoters (Fig. 2). The results indicate that transition amplitudes for RNAP-dependent DNA unwinding upon formation of RPo with CCT are larger than those for formation of RPo with GGG, on both positively and negatively supercoiled templates (Fig. 2B, left and center). Transition-amplitude histograms confirm that transition amplitudes with CCT are larger than with GGG, on both positively and negatively supercoiled templates (Fig. 2B, right). By combining the results with positively and negatively supercoiled templates to deconvolve effects of RNAP-dependent DNA unwinding and RNAP-dependent compaction (Revyakin et al., 2004, 2005, 2006), we find a 2 bp difference in RNAP-dependent DNA unwinding for CCT vs. GGG (Fig. 2C), corresponding exactly to the 2 bp difference in TSS selection (Fig. 1B). We conclude that TSS selection shows the second hallmark of scrunching.

### TSS selection downstream and upstream of modal TSS involves scrunching and anti-scrunching, respectively: forward and reverse movements of RNAP leading edge

According to the hypothesis that TSS selection involves scrunching or anti-scrunching, TSS selection at the most frequently observed, modal TSS position (7 bp downstream of −10 element for majority of discriminator sequences, including GGG) involves neither scrunching nor anti-scrunching, TSS selection downstream of the modal position involves scrunching (transcription-bubble expansion), and TSS selection upstream of the modal position involves anti-scrunching (transcription-bubble contraction; Robb et al., 2013; Vvedenskaya et al., 2015; Winkelman et al., 2016a,b). The results in Figs. 1–2 apply to the modal TSS position and a TSS position 2 bp downstream of the modal TSS position. To generalize and extend the results to a range of different TSS positions, including a position upstream of the modal TSS position expected to involve anti-scrunching, we exploited the ability of oligoribonucleotide primers (“nanoRNAs”) (Goldman et al., 2011) to program TSS selection (Figs. 3A, S3). We analyzed a consensus bacterial promoter, lacCONS, and used four ribotrinucleotide primers, UGG, GGA, GAA, and AAU, to program TSS selection at positions 6, 7, 8, and 9 bp downstream of the − 10 element (Fig. 3A). Experiments analogous to those in Fig. 1 show a one-for-one, bp-for-bp correlation between primer-programmed changes in TSS and changes in RNAP-leading-edge position. The leading-edge crosslink positions with the four primers differed in single-nucleotide increments, but the trailing-edge crosslink positions were the same (Fig. 3B). With the primer GGA, which programs TSS selection at the modal position (7 bp downstream of −10 element for this discriminator sequence), the leading-edge crosslinks were exactly as in experiments with no primer (Fig. S3). With primers GAA and AAU, which program TSS selection 1 and 2 bp downstream (positions 8 and 9), leading-edge crosslinks were 1 and 2 bp downstream of crosslinks with GGA (Fig. 3B). With primer UGG, which programs TSS selection 1 bp upstream (position 6), leading-edge crosslinks were 1 bp upstream of crosslinks with GGA (Fig. 3B). The results show that successive single-bp changes in TSS selection are matched by successive single-bp changes in RNAP leading-edge position. We conclude that the first hallmark of scrunching is observed for a full range of TSS positions, including, importantly, a position upstream of the modal TSS expected to involve anti-scrunching.

### TSS selection downstream and upstream of modal TSS involves scrunching and anti-scrunching, respectively: increases and decreases in RNAP-dependent DNA unwinding

We next used magnetic-tweezers single-molecule DNA-nanomanipulation to analyze primer-programmed TSS selection. To enable single-bp resolution, we reduced the DNA-tether length from 2.0 kb to 1.3 kb, thereby reducing noise due to compliance (Fig. S4; see Revyakin et al., 2005). The resulting transition amplitudes, transition-amplitude histograms, and RNAP-dependent DNA unwinding values for TSS selection with the four primers show a one-for-one, bp-for-bp correlation between primer-programmed changes in TSS and changes in RNAP-dependent DNA unwinding (Fig. 4). With primer GGA, which programs TSS selection at the modal position (7 bp downstream of the −10 element for this discriminator sequence), DNA unwinding was exactly as in experiments with no primer (Fig. S5). With primers GAA and AAU, which program TSS selection 1 and 2 bp further downstream (positions 8 and 9), DNA unwinding was ~1 and ~2 bp greater than with GGA (Fig. 4). With primer UGG, which programs TSS selection 1 bp upstream, DNA unwinding was ~1 bp less than that in experiments with GGA (Fig. 4). The results show that successive single-base-pair changes in TSS selection are matched by successive single-base-pair changes in DNA unwinding for a full range of TSS positions including, importantly, a position upstream of the modal TSS expected to involve anti-scrunching. Taken together, the results of protein-DNA photocrosslinking (Figs. 3, S3) and DNA-nanomanipulation (Figs. 4, S5) demonstrate, definitively, the scrunching/anti-scrunching hypothesis for TSS selection.

### Energetic costs of scrunching and anti-scrunching

To quantify the energetic costs of scrunching and anti-scrunching, we measured primer-concentration dependences of lifetimes of unwound states (Figs. 5–6, S6). For each primer, increasing the primer concentration increases the lifetime of the unwound state (t_unwound_), as expected for coupled equilibria of promoter unwinding, promoter scrunching, and primer binding (Figs. 5–6). The results in Fig. 6D show that the slopes of plots of mean t_unwoun_ (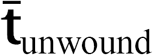) vs. primer concentration differ for different primers. Fitting the results to the equation describing the coupled equilibria (Fig. 6C) yields values of K_NpNpN_, ΔG_NpNpN_, K_scrunch_, and ΔG_scrunch_ for the four primers (Figs. 6E, S6). The results indicate that scrunching by 1 bp requires 0.8 kcal/mol, scrunching by 2 bp requires 2.0 kcal/mol, and anti-scrunching by 1 bp requires 1.5 kcal/mol (Figs. 6E, S6). The results provide the first experimental determination of the energetic costs of scrunching and anti-scrunching in any context.

### Energetics costs of scrunching and anti-scrunching in TSS selection explain range and relative utilization of TSS positions

The ΔG_scrunch_ values for bacterial TSS selection obtained in this work account for the range of TSS positions and the relative utilization of different TSS positions in bacterial transcription initiation. The ΔG_scrunch_ values for TSS selection at positions 6, 7, 8, and 9 of a promoter where the modal TSS is position 7 (0-2 kcal/mol) all are less than or comparable to 3*k*_B_T (~2 kcal/mol), where *k*_B_ is the Bolztmann constant and T is temperature in °K, indicating that TSS selection at these positions requires no energy beyond energy available in the thermal bath. Indeed, the probabilities of TSS selection at positions 6, 7, 8, and 9 as observed in a comprehensive analysis of TSS-region sequences (8%, 55%, 29%, 7%) (Vvedenskaya et al., 2015) can be predicted from the Boltzmann-distribution probabilities for the ΔG_scrunch_ values for TSS selection at these positions (6%, 72%, 18%, 3%; Fig. 6F). The finding that values of ΔG_scrunch_ for scrunching and anti-scrunching in TSS selection are ~1 kcal mol^-1^ bp^-1^ and 1-2 kcal mol^-1^ bp^-1^, respectively, implies that TSS selection at positions >2 bp downstream or >1 bp upstream of the modal position would exceed the energy fluctuations available to 99% of molecules at 20-37°C, and therefore explains the observation that TSS selection >2 bp downstream or >1 bp upstream of the modal position occurs rarely (Vvedenskaya et al., 2015).

### Unified mechanism of TSS selection by multisubunit RNAP

TSS selection by archaeal RNAP, eukaryotic RNAP I, eukaryotic RNAP II from most species, and eukaryotic RNAP III involves the same range of TSS positions as TSS selection by bacterial RNAP (positions ±2 bp from the modal TSS; Learned and Tjian, 1982; Samuels et al., 1984; Thomm and Wich, 1988; Reiter et al., 1990; Fruscoloni et al., 1995; Zecherle et al., 1996). We propose that TSS selection by all of these enzymes is mediated by scrunching and anti-scrunching driven by energy available in the thermal bath. In contrast, TSS selection by *S. cerevisiae* RNAP II involves a range of TSS positions of 10s to 100s of bp (long-range TSS scanning) (Giardina and Lis, 1993; Kuehner and Brow, 2006). We propose that TSS scanning by *S. cerevisiae* RNAP II also is mediated by scrunching and anti-scrunching, but, in this case, involves not only energy from the thermal bath, but also energy from the ATPase activity of RNAP II transcription factor TFIIH (Sainsbury et al., 2015). This proposal could account for the ATP-dependent, TFIIH-dependent cycles of DNA compaction and de-compaction of 10s to 100s of bp observed in single-molecule optical-tweezer analyses of TSS scanning by *S. cerevisiae* RNAP II (Fazal et al., 2015).

## ACKNOWLEDGEMENTS

Work was supported by NIH grants GM041376 (R.H.E), GM118059 (B.E.N), and a European Science Foundation EURYI grant (T.R.S). We thank Seth Goldman for help with construction of pCDF-CP. T.R.S, B.E.N, R.H.E designed and supervised the experiments.

## AUTHOR CONTRIBUTIONS

L.Y. performed DNA-nanomanipulation experiments. J.T.W. and C.P. performed protein-DNA crosslinking experiments. L.Y., J.T.W., C.P., T.R.S., B.E.N, and R.H.E designed experiments and analyzed data. L.Y., J.T.W., B.E.N., and R.H.E. wrote the paper. The authors declare that no competing interests exist.

---

**Fig. 1.**
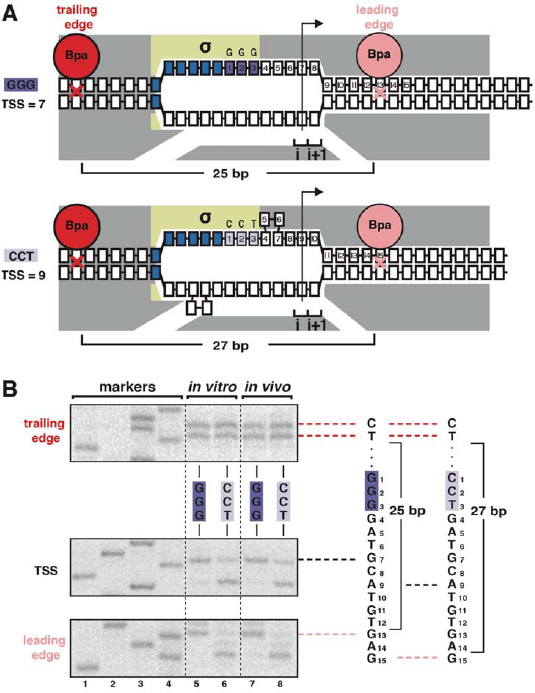
TSS selection exhibits first hallmark of scrunching--movement of RNAP leading edge but not RNAP trailing edge--both *in vitro* and *in vivo.* (**A**)RNAP leading-edge and trailing-edge positions at promoters having GGG discriminator (TSS 7 bp downstream of −10 element; top) and CCT discriminator (TSS 9 bp downstream of −10 element; bottom). Changes in TSS selection result from changes in discriminator-sequence-dependent DNA scrunching. Gray, RNAP; yellow, c; blue, −10-element nucleotides; dark purple, GGG-discriminator nucleotides; light purple, CCT-discriminator nucleotides; i and i+1, NTP binding sites; arrow, TSS; boxes, DNA nucleotides (nontemplate-strand nucleotides above template-strand nucleotides; nucleotides downstream of −10 element numbered); red, trailing-edge Bpa and nucleotide crosslinked to Bpa; pink, leading-edge Bpa and nucleotide crosslinked to Bpa. Scrunching is indicated by bulged-out nucleotides. Distance between leading-edge and trailing-edge crosslinks is indicated below RNAP. (**B**) RNAP trailing-edge crosslinking (top), TSS (middle), and RNAP leading-edge crosslinking (bottom) for promoters having GGG discriminator and CCT discriminator, *in vitro* (lanes 5-6) and *in vivo* (lanes 7-8). Horizontal dashed lines relate bands on gel (left) to nucleotide sequences (right).

**Fig. 2.**
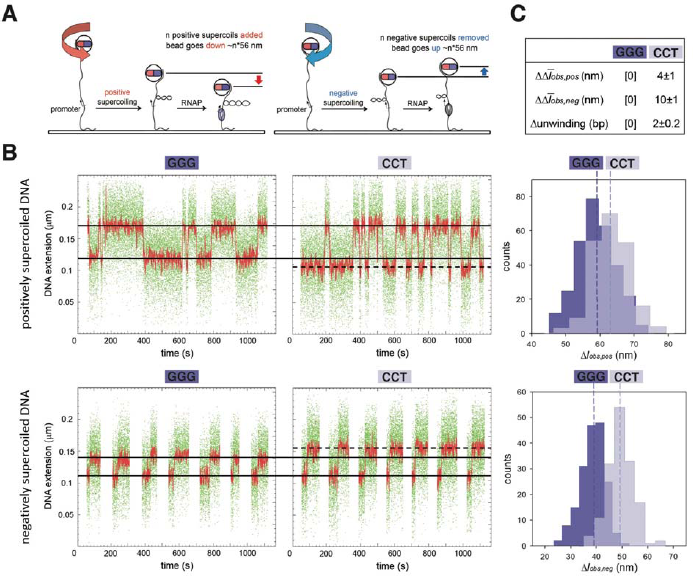
TSS selection exhibits second hallmark of scrunching--change in size of transcription bubble. (**A**) Magnetic-tweezers single-molecule DNA nanomanipulation (see Revyakin et al., 2004, 2005, 2006). End-to-end extension (*l*) of a mechanically stretched, positively supercoiled (left), or negatively supercoiled (right), DNA molecule is monitored. Unwinding of *n* turns of DNA by RNAP results in compensatory gain of *n* positive supercoils or loss of *n* negative supercoils, and movement of the bead by *n*\*56 nm. (**B**) Single-molecule time traces and transition-amplitude histograms for RPo at promoters having GGG discriminator or CCT discriminator. Upper subpanel, positively supercoiled DNA; lower subpanel, negatively supercoiled DNA. Green points, raw data (30 frames/s); red points, averaged data (1-s window); horizontal black lines, wound and unwound states of GGG promoter; dashed horizontal black line, unwound state of CCT promoter; vertical dashed lines, means; Δ*l*_*obs,pos,*_ transition amplitude with positively supercoiled DNA; *Al_obs_,_neg_,* transition amplitude with negatively supercoiled DNA; 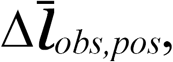 mean Δ*l*_*obs, pos*_; 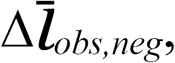 mean Δ*l*_*obs, neg*_. (**C**) Differences in 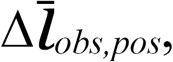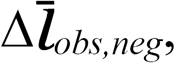 and DNA unwinding between GGG-discriminator promoter and CCT-discriminator promoter (means±SEM).

**Fig. 3.**
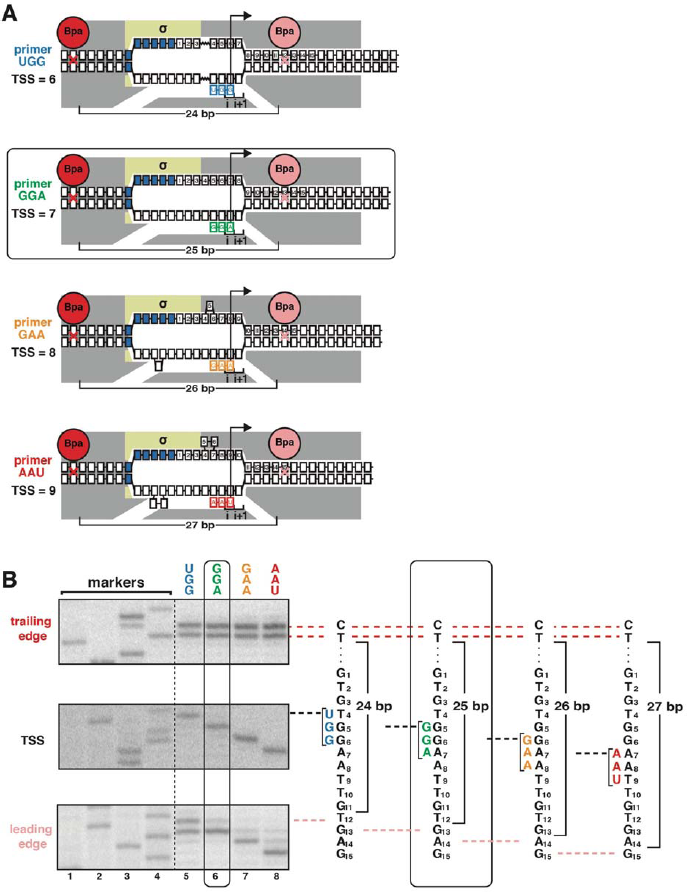
TSS selection downstream and upstream of the modal TSS involves, respectively: forward and reverse movements of RNAP leading edge. (**A**) Ribotrinucleotide primers program TSS selection at positions 6, 7, 8, and 9 bp downstream of −10 element (UGG, GGA, GAA, and AAU). Cyan, green, orange, and red denote primers UGG, GGA, GAA, and AAU, respectively. Rectangle with rounded corners highlights case of primer GGA, which programs TSS selection at same position as in absence of primer (7 bp downstream of −10 element). Other colors as in Fig. 1A. (**B**) Use of protein-DNA photocrosslinking to define RNAP leading-edge and trailing-edge positions. RNAP trailing-edge crosslinking (top), TSS (middle), and RNAP leading-edge crosslinking (bottom) with primers UGG, GGA, GAA, and AAU (lanes 5-8). Horizontal dashed lines relate bands on gel (left) to nucleotide sequences (right).

**Fig. 4.**
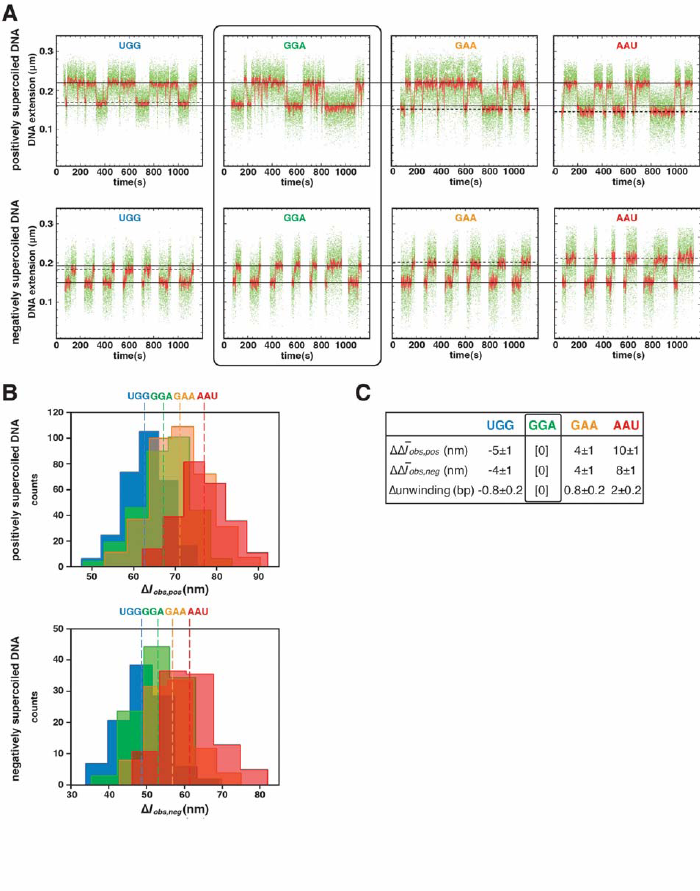
TSS selection downstream and upstream of the modal TSS involves, respectively: increases and decreases in RNAP-dependent DNA unwinding. (**A**) Use of single-molecule DNA nanomanipulation to define RNAP-dependent DNA unwinding. Single-molecule time traces with primers UGG, GGA, GAA, and AAU. Rectangle with rounded corners highlights case of primer GGA, which programs TSS selection at position 7. Colors as in Fig. 2B. (**B**) Transition-amplitude histograms (positively supercoiled DNA in top panel; negatively supercoiled DNA in middle panel). (**C**) Differences in 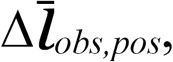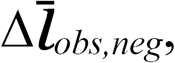 and DNA unwinding (bottom panel) with primers UGG, GGA, GAA, and AAU (means±SEM).

**Fig. 5.**
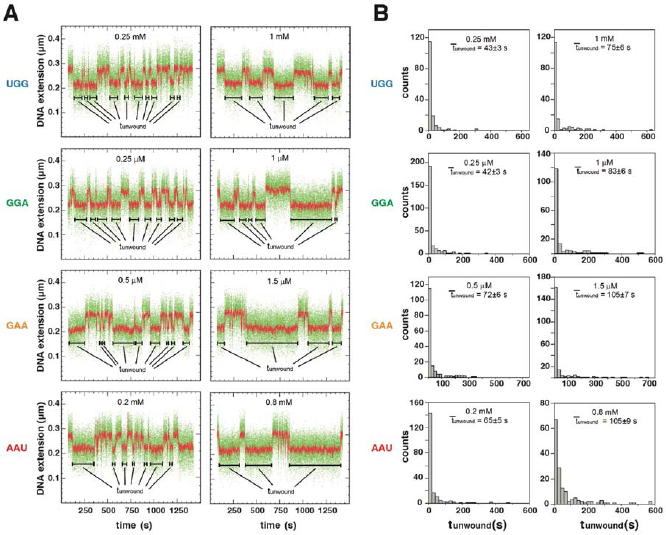
Energetic costs of scrunching and anti-scrunching: primer-concentration dependences of unwound-state lifetimes with primers UGG, GGA, GAA, and AAU. (**A**) Single-molecule time traces at low (left) and high (right) primer concentrations. Black bars, lifetimes of unwound states (t_unwound_). Colors as in Fig. 2B. (**B**) tunwound distributions and mean tunwound (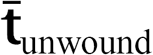). 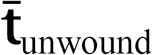 increases with increasing primer concentrations.

**Fig. 6.**
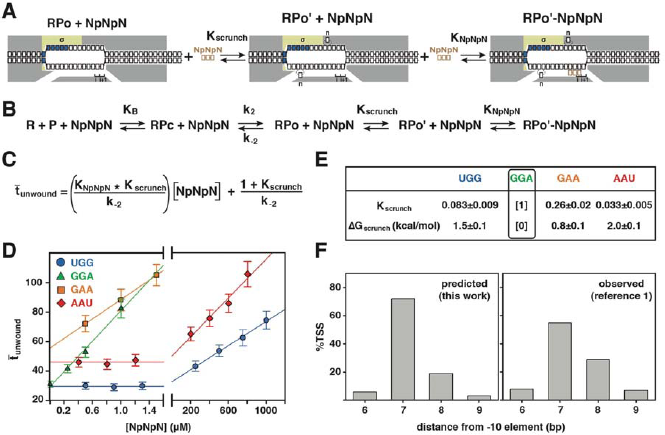
Energetic costs of scrunching and anti-scrunching: calculation of energetic costs and relationship between energetic costs and range and relative utilization of TSS positions. (**A**) Coupled equilibria for promoter scrunching or anti-scrunching (K_scrunch_) and primer binding (K_NpNpN_). (**B**) Equations for promoter binding (K_B_), promoter unwinding (k_2_/k_2_), promoter scrunching or anti-scrunching (K_scrunch_), and primer binding (K_NPNPN_). (**C**) Relationship between 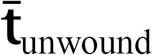, K_scrunch_, K_NpNpN_, and primer concentration. # (**D**) Dependences of mean lifetimes of unwound states (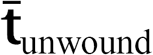) on primer concentration for primers UGG, GGA, GAA, and AAU (means±SEM). (**E**) Values of K_scrunch_ and ΔG_scrunch_ calculated by fitting data in **D** to equation in **C**, using 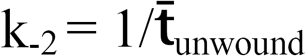 in absence of primer (Revyakin et al., 2004) (means±SEM). Colors as in Fig. 3. (**F**) TSS distributions predicted by Boltzmann-distribution probabilities for ΔG_scrunch_ values in E (left) and observed in analysis of comprehensive library of TSS-region sequences in Vvedenskaya et al., 2015) (right).

## METHODS

### Proteins

Wild-type *E. coli* RNAP core enzyme used in transcription experiments was prepared from *E. coli* strain NiCo21(DE3) (New England BioLabs) transformed with plasmid pIA900 (Svetlov and Artsimovitch, 2015) as described (Winkelman et al., 2015). Wild-type RNAP for single-molecule DNA-nanomanipulation experiments was prepared from *E. coli* strain BL21(DE3) (New England Biolabs) transformed with plasmid pVS10 (Artsimovitch et al., 2003) as in Artsimovitch et al., 2003.

Bpa-containing RNAP core-enzyme derivatives for *in vitro* protein-DNA photocrosslinking (β’R1148Bpa for analysis of RNAP leading-edge positions; β′T48Bpa for analysis of RNAP trailing-edge positions) were prepared from *E. coli* strain NiCo21(DE3) (New England BioLabs) co-transformed with plasmid pEVOL-pBpF (Chin et at., 2002; Addgene) and plasmid pIA900-β′R1148Bpa (Winkelman et al., 2015) or pIA900-β′T48Bpa (Winkelman et al., 2015), as in Winkelman et al., 2015.

Bpa-containing, mutationally inactivated, RNAP core-enzyme derivatives for *in vitro* and *in vivo* protein-DNA photocrosslinking (β′R1148Bpa;β′D460A for analysis of RNAP leading-edge positions; β′T48Bpa;β′D460A for analysis of RNAP trailing-edge positions) were prepared from *E. coli* strain NiCo21(DE3) (New England BioLabs) co-transformed with plasmid pEVOL-pBpF (Chin et at., 2002; Addgene) and plasmid pIA900-β′R1148Bpa;β′D460A or pIA900-β′T48Bpa;β′D460A [constructed from pIA900-β′R1148-Bpa (Winkelman et al., 2015) and pIA900-β′T48-Bpa (Winkelman et al., 2015) by use of site-directed mutagenesis with primer "JW30", as in Winkelman et al., 2015].

σ^70^ was prepared from *E. coli* strain BL21(DE3) (New England Biolabs) transformed with plasmid pσ^70^-His (Marr and Roberts, 1997) as in Marr and Roberts, 1997. To form RNAP holoenzyme, 1 μM RNAP core enzyme and 5 μM σ^70^ in 10 mM Tris-Cl, pH 8.0, 100 mM KCl, 10 mM MgCl_2_, 0.1 mM EDTA, 1 mM DTT, and 50% glycerol were incubated 30 min at 25°C.

### Oligonucleotides

Oligodeoxyribonucleotides (desalted) were purchased from IDT (sequences in Table S1). Oligoribonucleotides (HPLC-purified) were purchased from Trilink Biotechnologies.

### Determination of TSS position *in vitro*

Experiments in Figs. 1B and S2B were performed using reaction mixtures (60 μ1) containing 20 nM RNAP holoenzyme derivative, 4 nM plasmid pCDF-CP-*lac*CONS-GGG or pCDF-CP-*lac*CONS-CCT, carrying derivatives of the *lac*CONS promoter (Mukhopadhyay et al., 2001) having a GGG or CCT discriminator [prepared by inserting a synthetic 248 bp DNA fragment (5′-GAAGCCCTGCATTAGGGGTACCCTAGAGCCTGACCGGCATTATAGCCCCAGCGGCGGATCCC TGCGGGTCGACAAGCTTGAATAGCCATCCCAATCGAACAGGCCTGCTGGTAATCGCAGGCCT TTTTATTTGGATGGAGCTCTGAGAGTCTTCGGTGTATGGGTTTTGCGGTGGAAACACAGAAA AAAGCCCGCACCTGACAGTGCGGGCTTTTTTTTTCGACCAAAGGGACGACCGGGTCGTTGGT − 3′) between positions 3601 and 460 of pCDF-1b (EMD-Millipore), yielding plasmid pCDF-CP, followed by ligating a 200 bp BglI-digested DNA fragment carrying the *lac*CONS promoter with GGG or CCT discriminator (5′-GTTCAGAGTTCTACAGTCCGACGATCGCGGATGCTTGACAGAGTGAGCGCAACGCAATAACAGTCATCTAGATAGAACTTTAGGCACCCCAGGCTTGACACTTTATGCTTCGGCTCGTATAAT **<u>GGG</u>**GATGCATGTGAGCGGATAACAATGCGGTTAGGCTTAGAGCGCTTAGTCGATGCTGGAA TTCTCGGGTGCCAAGG −3′ or 5′-GTTCAGAGTTCTACAGTCCGACGATCGCGGATGCTTGACAGAGTGAGCGCAACGCAATAACAGTCATCTAGATAGAACTTTAGGCACCCCAGGCTTGACACTTTATGCTTCGGCTCGTATAAT **CCT**GATGCATGTGAGCGGATAACAATGCGGTTAGGCTTAGAGCGCTTAGTCGATGCTGGAA TTCTCGGGTGCCAAGG −3′; −35 and −10 elements underlined; discriminator in bold) with BglI-digested plasmid pCDF-CP], 0 or 1 mM ATP, 0 or 1 mM CTP, 0 or 1 mM GTP, and 0 or 1 mM UTP in 60 μl 10 mM Tris-Cl, pH 8.0, 70 mM NaCl, 10 mM MgCl_2_, and 0.1 mg/ml bovine serum albumin. After 20 min at 37°C, reactions were terminated by addition of 100 μl 10 mM EDTA pH 8.0 and 1 mg/ml glycogen. Samples were extracted with acid phenol:chloroform (Sambrook and Russell, 2001) (Ambion), and RNA products were recovered by ethanol precipitation (Sambrook and Russell, 2001) and re-suspended in 6.5 μl water. The RNA products were analyzed by primer extension to define TSS positions. Primer-extension was performed by combining 6.5 μl RNA products in water, 1 μl 1 μM ^32^P-5′-end-labeled primer "s128a" [Table S1; 200 Bq/fmol; prepared using [γ^32^P]-ATP (PerkinElmer) and T4 polynucleotide kinase (New England Biolabs); procedures as in Sambrook and Russel, 2001], and 1 μl 10x avian myelobastosis virus (AMV) reverse transcriptase buffer (New England BioLabs) heating 10 min at 90, cooling to 40°C at 0.1°C/s, and incubating 15 min at 40°C; adding 0.5 μl 10 mM dNTP mix (2.5 mM dATP, 2.5 mM dGTP, 2.5 mM, dCTP, and 2.5 mM dTTP; New England Biolabs) and 1 μl 10 U/μl AMV reverse transcriptase (New England BioLabs); and incubating 1 h at 50°C. Primer-extension reactions were terminated by heating 20 min at 85°C; 10 μl 1x TBE (Artsimovitch et al., 2003), 8 M urea, 0.025% xylene cyanol, and 0.025% bromophenol blue was added; and samples were analyzed by electrophoresis on 8 M urea, 1X TBE polyacrylamide gels UreaGel System; National Diagnostics) (procedures as in Sambrook and Russel, 2001), followed by storage-phosphor imaging (Typhoon 9400 variable-mode imager; GE Life Science). TSS positions were determined by comparison to products of a DNA-nucleotide sequencing reaction obtained using a PCR-generated DNA fragment containing positions −129 to +71 of the *lac*CONS-GGG promoter and primer "s128a" (Thermo Sequenase Cycle Sequencing Kit; Affymetrix; methods as per manufacturer). Experiments in Fig. 3B, were performed analogously, but using a 1.3 kb DNA fragment carrying positions −687 to +644 of the *lac*CONS promoter (Mukhopadhyay et al., 2001) prepared by PCR amplification of plasmid pUC18-T20C2-lacCOAS [prepared by replacing the SbfI-XbaI segment of plasmid pUC18 (Thermo Scientific) with a 2.0 kb Sbfl-Xbal DNA fragment obtained by PCR amplification of *Thermus aquaticus rpoC* gene with primers Taq_rpoC_F and Taq_rpoC_R (Table S1) and digestion with XbaI and SbfI-HF (New England BioLabs), yielding plasmid pUC18-T20C2, followed by inserting a synthetic 117 bp DNA fragment carrying the lacCONS promoter (5′-CGGATGCTTGACAGAGTGAGCGCAACGCAATAACAGTCATCTAGATAGAACTTTAGGCACC CCAGGCTTGACACTTTATGCTTCGGCTCGTATAAT*GTG*TGGAATTGTGAGCGGATA-3′; −35 and −10 elements underlined; discriminator in bold) into the KpnI site of plasmid pUC18-T20C2] with primers “LY10” and "LY11" (Table S1), and performing experiments in the presence of 0 or 1 mM UGG, GGA, GAA, or AAU.

### Determination of TSS position *in vivo*

*E. coli* strain NiCo21(DE3) (New England BioLabs) transformed with plasmid pCDF-CP-lacCONS-GGG or pCDF-CP-lacCONS-CCT was plated on LB agar (Sambrook and Russell, 2001) containing 50 μg/ml spectinomycin and 50 μg/ml streptomycin, single colonies were inoculated into 25 ml LB broth (Artsimovitch et al., 2003) containing 50 μg/ml spectinomycin and 50 μg/ml streptomycin in 125 ml Bellco flasks, and cultures were shaken (220 rpm) at 37°C. When cell densities reached OD_600_ = 0.6, 2 ml aliquots were centrifuged 2 min at 4°C at 23,000xg, and resulting cell pellets were frozen at −80°C. Cell pellets were thawed in 1 ml TRI Reagent (Molecular Research Center) at 25°C for 5 min, completely re-suspended by pipetting up and down, incubated 10 min at 70°C, and centrifuged 2 min at 25°C at 23,000 x g. Supernatants were transferred to fresh 1.7 ml microfuge tubes, 200 μl chloroform (Ambion) was added, vortexed, and samples were centrifuged 1 min at 25°C at 23,000 x g. Aqueous phases were transferred to a fresh tube and nucleic acids were extracted with acid phenol:chloroform (Sambrook and Russell, 2001). Nucleic acids were recovered by ethanol precipitation (Sambrook and Russell, 2001), and re-suspended in 20 μl 10 mM Tris-Cl, pH 8.0. Primer extension was performed as described in the preceding section.

### Determination of RNAP leading-edge and trailing-edge positions *in vitro:* protein-DNA photocrosslinking *in vitro*

Experiments in Fig. 1B were performed using reaction mixtures (10 μl) containing 50 nM Bpa-containing RNAP holoenzyme derivative β′R1148Bpa (for analysis of RNAP leading-edge positions) or β′T48Bpa (for analysis of RNAP trailing-edge positions) and 4 nM plasmid pCDF-CP-*lac*CONS-GGG or plasmid pCDF-CP-*lac*CONS-CCT in 10 mM Tris-Cl, pH 8.0, 70 mM NaCl, 10 mM MgCl2, and 0.1 mg/ml bovine serum albumin. Reaction mixtures were incubated 5 min at 37°C, UV-irradiated 5 min at 25°C in a Rayonet RPR-100 photochemical reactor equipped with 16 x 350 nm tubes (Southern New England Ultraviolet), and resulting protein-DNA crosslinks were mapped using primer extension. Primer-extension reactions (12.5 μl) were performed by combining 2 μl aliquot of crosslinking reaction, 1 μl 1 μM ^32^P-5′-end-labeled primer "s128a" (for analysis of leading-edge positions) or primer "JW85" (for analysis of trailing-edge positions) [Table S1; 200 Bq/fmol; prepared using [γ^32^P]-ATP (PerkinElmer) and T4 polynucleotide kinase (New England Biolabs); procedures as in Sambrook and Russell, 2001], 1 μl 10X dNTPs (2.5 mM dATP, 2.5 mM dCTP, 2.5 mM dGTP, 2.5 mM TTP, 0.5 μl 5 U/μl Taq DNA polymerase (New England BioLabs), 5 μl 5 M betaine, 0.625 μl 100% dimethyl sulfoxide, and 1.25 μl 10x Taq DNA polymerase buffer (New England BioLabs); and cycling 16-40 times through 30 s at 95°C, 30 s at 53°C, and 30 s at 72°C. Primer-extension reactions were terminated, and primer-extension products were analyzed as in the preceding section. Experiments in Fig. S2B were performed analogously, but using Bpa-containing, mutationally inactivated, RNAP derivatives β′R1148Bpa; β′D460A (for analysis of RNAP leading-edge positions) and β′T48Bpa; β′D460A (for analysis of RNAP trailing-edge positions)

Experiments in Figs. 3B and S3B were performed analogously, but using reaction mixtures also containing 0 or 1 mM of ribotrinucleotide primers UGG, GGA, GAA, or AAU, and using ^32^P-5′-end-labeled primers "JW62" and "JW61" (Table S1) in primer-extension reactions.

### Determination of RNAP leading-edge and trailing-edge positions *in vivo:* protein-DNA photocrosslinking *in vivo*

Experiments in Fig. 1B were performed using a three-plasmid merodiploid system that enabled production of a Bpa-containing, mutationally inactivated, decahistidine-tagged RNAP holoenzyme derivative *in vivo* and enabled trapping of RPo consisting of the Bpa-containing, mutationally inactivated, decahistidine-tagged RNAP holoenzyme derivative and a *lac*CONS promoter with GGG or CCT discriminator *in vivo,* and UV-irradiation of cells (Fig. S1).

*E. coli* strain NiCo21(DE3) (New England BioLabs) transformed sequentially with (1) plasmid pCDF-CP-*lac*CONS-GGG or plasmid pCDF-CP-*lac*CONS-CCT, (2) plasmid pIA900-β′T48Bpa; β′D460A or plasmid pIA900-β′R1148Bpa; β′D460A, and (3) plasmid pEVOL-pBpF (Chin et al., 2002; Addgene) was plated to yield a confluent lawn on LB agar (Sambrook and Russell, 2001) containing 100 μg/ml carbenicillin, 50 μg/ml spectinomycin, 50 μg/ml streptomycin, and 25 μg/ml chloramphenicol; cells were scraped from the plate and used to inoculate 250 ml LB broth (as described above) containing 1 mM Bpa (Bachem), 100 μg/ml carbenicillin, 50 μg/ml spectinomycin, 50 μg/ml streptomycin, and 25 μg/ml chloramphenicol in a 1000 ml flask (Bellco) to yield OD_600_ = 0.3; the culture was shaken (220 rpm) 1 h at 37°C in the dark, isopropyl-β-D-thiogalactoside was added to 1 mM; and the culture was further shaken (220 rpm) 3 h at 37°C in the dark. Aliquots (7 ml) were transferred to 13 mm x 100 mm borosilicate glass test tubes (VWR), UV-irradiated 20 min at 25°C in a Rayonet RPR-100 photochemical reactor equipped with 16 x 350 nm tubes (Southern New England Ultraviolet), harvested by centrifuging 15 min at 4°C at 3000xg, and cell pellets were frozen at −20°C. Cell pellets were thawed 30 min at 4°C, re-suspended in 40 ml 50 mM Na_2_HPO_4_ pH 8.0, 1.4 M NaCl, 20 mM imidazole, 14 mM β-mercaptoethanol, 0.1% Tween20, and 5% ethanol containing 2 mg egg white lysozyme. Cells were lysed by sonication 5 min at 4°C., cell lysates were centrifuged 40 min at 4°C at 23,000xg, and supernatants were added to 1 ml Ni-NTA-agarose (Qiagen) in 1 ml 50 mM Na_2_HPO_4_, pH 8.0, 1.4 M NaCl, 20 mM imidazole, 0.1% Tween-20, 5 mM β-mercaptoethanol, and 5% ethanol, and incubated 30 min at 4°C with gentle rocking. The Ni-NTA-agarose was loaded into a 15 ml polyprep column (BioRad), the resulting column was washed with 10 ml of 50 mM Na_2_HPO_4_, pH 8.0, 300 mM NaCl, 20 mM imidazole, 0.1% Tween-20, 5 mM β-mercaptoethanol, and 5% ethanol and eluted with 3 ml of the same buffer containing 300 mM imidazole. The eluate was concentrated to 0.2 ml using an 1000 MWCO Amicon Ultra-4 centrifugal filter (EMD Millipore); the buffer was exchanged to 0.2 ml 20 mM Tris-Cl, pH 8.0, 200 mM KCl, 20 mM MgCl_2_, 0.2 mM EDTA, and 1 mM DTT using the 1000 MWCO Amicon Ultra-4 centrifugal filter (EMD Millipore); 0.2 ml glycerol was added; and the sample was stored at −20°C. Protein-DNA crosslinks were mapped by denaturation followed by primer extension. Denaturation was performed by combining 25 μl crosslinked RNAP-DNA, 25 μl water, 15 μl 5 M NaCl, and 6 μl 100 μg/ml heparin; heating 5 min at 95°C; cooling on ice. Denatured crosslinked RNAP-DNA was purified by adding 20 μl MagneHis Ni-particles (Promega) equilibrated and suspended in 10 mM Tris-Cl, pH 8.0, 1.2 M NaCl, 10 mM MgCl_2_, 10 μg/ml heparin, and 0.1 mg/ml bovine serum albumin; washing once with 50 μl 10 mM Tris-Cl, pH 8.0, 1.2 M NaCl, 10 mM MgCl_2_, 10 μg/ml heparin, and 0.1 mg/ml bovine serum albumin; washed twice with 50 μl 1x *Taq* DNA polymerase buffer (New England BioLabs); and resuspended in 10 μl 1 × *Taq* DNA polymerase buffer. Primer extension was performed using 2 μl aliquots of purified denatured crosslinked RNAP-DNA, using procedures essentially as described above for experiments in Fig. 1B.

### Determination of RNAP-dependent DNA unwinding by single-molecule DNA-nanomanipulation: DNA constructs

2.0 kb DNA fragments carrying single centrally located *lac*CONS-GGG, *lac*CONS-CCT, or **lac*CONS* promoters were prepared by digesting plasmid pUC18-T20C2-*lac*CONS-GGG or plasmid pUC18-T20C2-*lac*CONS-CCT [prepared by inserting a synthetic 80 bp DNA fragment carrying a derivative of the *lac*CONS promoter (Mukhopadhyay et al., 2001) having a GGG or CCT discriminator (5′-CATCTAGATCACATTTTAGGCACCCCAGGCTTGACACTTTATGCTTCGGCTCGTATAAT**GGG**GATGCATGTGAGCGGATA-3′ or 53′-CATCTAGATCACATTTTAGGCACCCCAGGCTTGACACTTTATGCTTCGGCTCGTATAAT**CCTG**ATGCATGTGAGCGGATA −3′; −35 and −10 elements underlined; discriminator in bold) into the KpnI site of plasmid pUC18-T20C2] or plasmid pUC18-T20C2-*lac*CONS with XbaI and SbfI-HF (New England BioLabs), followed by agarose gel electrophoresis.

1.3 kb DNA fragments carrying a single centrally located **lac*CONS* promoter were prepared by PCR amplification of plasmid pUC18-T20C2-*lac*CONS, using primers "LY10" and "LY11" (Table S1), followed by treatment with Dam methyltransferase (New England BioLabs), digestion with XbaI and SbfI-HF (New England BioLabs), and agarose gel electrophoresis.

DNA constructs for magnetic-tweezers single-molecule DNA-nanomanipulation were prepared from the above 2.0 kb and 1.3 kb DNA fragments by ligating, at the XbaI end, a 1.0 kb DNA fragment bearing multiple biotin residues on both strands [prepared by PCR amplification of plasmid pARTaqRPOC-*lac*CONS using primers "XbaRPOC4050" and "RPOC3140" (Table S1) and conditions as described (Revyakin et al., 2003, 2004, 2005, 2006), followed by digestion with XbaI (New England BioLabs) and agarose gel electrophoresis] and, at the SbfI end, a 1.0 kb DNA fragment bearing multiple digoxigenin residues on both strands [prepared by PCR amplification of plasmid pARTaqRPOC-*lac*CONS using primers "SbfRPOC50" and "RPOC820" (Table S1) and conditions as described (Revyakin et al., 2003, 2004, 2005, 2006), followed by digestion with SbfI-HF (New England BioLabs) and agarose gel electrophoresis]. DNA constructs were attached to streptavidin-coated magnetic beads (MyOne Streptavidin C1, Life Technologies), attached to anti-digoxigenin-coated glass surfaces (Lionnet et al., 2012), and calibrated as described (Revyakin et al., 2003, 2004, 2005, 2006).

### Determination of RNAP-dependent DNA unwinding by single-molecule DNA-nanomanipulation: data collection

Experiments were performed essentially as described (Revyakin et al., 2003, 2004, 2005, 2006).

Experiments in Fig. 2 (experiments addressing TSS selection for promoters with GGG or CCT discriminator sequence), were performed using standard reactions containing mechanically extended, torsionally constrained, 2.0 kb DNA molecule carrying GGG or CCT promoter (extension force = 0.3 pN; superhelical density = 0.021 for experiments with positively supercoiled DNA; superhelical density = − 0.021 for experiments with negatively supercoiled DNA) and RNAP holoenzyme (10 nM for experiments with positively supercoiled DNA; 0.5 nM for experiments with negatively supercoiled DNA) in 25 mM Na-HEPES, pH 7.9, 75 mM NaCl, 10 mM MgCl_2_, 1 mM dithiothreitol, 0.1% Tween-20, 0.1 mg/ml bovine serum albumin) at 30°C. Data from each of three single DNA molecules were pooled [differences in plectoneme size (at superhelical density +/−0.021) and 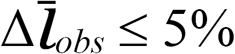].

Experiments in Fig. S4 (experiments demonstrating that reduction in DNA-fragment length from 2.0 to 1.3 kb enables single-bp resolution) were performed using standard reactions containing mechanically extended, torsionally constrained, 2.0 kb or 1.3 kb DNA molecule carrying *lac*CONS promoter (extension force = 0.3 pN; initial superhelical density = 0.021 or 0.0024 for experiments with 2.0 kb or 1.3 kb positively supercoiled DNA; superhelical density = −0.021 or −0.0024 for experiments with 2.0 kb or 1.3 kb negatively supercoiled DNA) and RNAP holoenzyme (10 nM for experiments with positively supercoiled DNA; 0.5 nM for experiments with negatively supercoiled DNA) in the buffer of the preceding paragraph at 30°C. For each DNA-fragment length, data were collected on one single DNA molecule.

Experiments in Figs. 4 and S5 (experiments addressing primer-programmed TSS selection with primers UGG, GGA, GAA, AAU) were performed using standard reactions containing mechanically extended, torsionally constrained 1.3 kb DNA molecule carrying *lac*CONS promoter (extension force = 0.3 pN; initial superhelical density = 0.024 for experiments with positively supercoiled DNA; superhelical density = −0.024 for experiments with negatively supercoiled DNA) and RNAP holoenzyme (10 nM for experiments with positively supercoiled DNA; 0.5 nM for experiments with negatively supercoiled DNA) in the buffer of the preceding paragraph at 30°C. Primers UGG, GGA, GAA, and AAU were present at 0 or 1 mM, 0 or 1 μM, 0 or 2.5 μM, and 0 or 1 mM, respectively. For experiments with positively supercoiled DNA, data from each of seven single DNA molecules were normalized based on 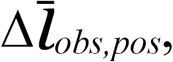 in absence of primer and pooled; for experiments with negatively supercoiled DNA, data from each of two single DNA molecules were normalized based on 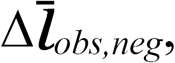 in absence of primer and pooled.

Experiments in Figs. 5–6 and S6 (experiments addressing primer-concentration dependences of t_unwound_ in primer-programmed TSS selection) were performed using standard reactions containing mechanically extended, torsionally constrained, 2.0 kb DNA molecule carrying *lac*CONS promoter (extension force = 0.3 pN; initial superhelical density = 0.021) and RNAP holoenzyme (10 nM) in the buffer of the preceding paragraph at 30°C. Each titration consisted of recordings in absence of primer followed by recordings in presence of primer at increasing concentrations. (0, 0.50, 0.90, 1.3, 250, 500, 750, and 1000 μM for UGG; 0, 0.25, 0.50, and 1.0 μM for GGA; 0, 0.50, 1.0, and 1.5 μM for GAA; 0, 0.40, 0.80, 1.2, 200, 400, 600, and 800 μM for AAU). For each titration, data were collected on one single DNA molecule.

For experiments with negatively supercoiled DNA, for which 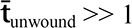 h (Revyakin et al., 2003, 2004, 2005, 2006), unwound DNA constructs were mechanically disrupted after 1 min essentially as described (Revyakin et al., 2003, 2004, 2005, 2006) [rotating magnets counterclockwise (8 turns or 6 turns for 2.0 kb DNA or 1.3 kb DNA) to introduce positive superhelical turns and disrupt unwound complexes (superhelical densiy = 0.021 or 0.024 for experiments with 2.0 kb or 1.3 kb DNA), waiting for ≥1 min, rotating magnets clockwise (8 turns or 6 turns for 2.0 kb DNA or 1.3 kb DNA) to re-introduce negative superhelical turns (superhelical density = −0.021 or −0.024 for 2.0 kb or 1.3 kb DNA) and allow re-formation of unwound complexes].

### Determination of RNAP-dependent DNA unwinding by single-molecule DNA-nanomanipulation: data reduction for determination of DNA unwinding

Raw time traces were analyzed to yield DNA extension (*l*) as described (Revyakin et al., 2003, 2004, 2005, 2006).

Changes in *l* attributable to DNA unwinding (Δ*l_μ_*) and changes in *l* attributable to DNA compaction (Δ*l_c_*) were calculated as: Δ*l_μ_* = (Δ*l_obs,neg_* + Δ*l_obs,pos_*)/2, and Δ*l_c_* = (Δ*l*_*obs,pos*_ − Δ*l*_*obs,neg*_)/2, where Δ*l*_*obs,pos*_ and Δ*l*_*obs,neg*_ are observed changes in *l* in experiments with positively supercoiled DNA and negatively supercoiled DNA, as described (Revyakin et al., 2003, 2004, 2005, 2006). Extents of DNA unwinding in base pairs were calculated as (Δ*l*_μ_/*δ*) 10.4, where *δ* (in the range of 50-65 nm) is the change in *l* per turn at superhelical densities used in this work (+/− 0.021 or +/− 0.024 for 2.0 kb DNA or 1.3 kb DNA), and 10.4 is the mean number of base pairs per turn of B-DNA (Wang, 1979), as described (Revyakin et al., 2003, 2004, 2005, 2006). Extents of DNA compaction in nm were calculated as (Δ*l*_*c*_)/0.7, where 0.7 is the fractional extension of DNA at 0.3 pN extension force (Bustamante et al., 1994), as described (Revyakin et al., 2003, 2004, 2005, 2006). Changes in DNA compaction associated with scrunching or anti-scrunching for these experiments are expected to be <1 nm for the experiments in Fig. 4 (0-2 bp times 0.34 nm/bp, where 0.34 nm/bp is the rise per base pair of B-DNA) (Franklin and Gosling, 1953); changes in DNA compaction on this scale are comparable to the error of this experimental approach and thus cannot be detected by this approach (Revyakin et al., 2003, 2004, 2005, 2006).

### Determination of RNAP-dependent DNA unwinding by single-molecule DNA-nanomanipulation: data reduction for determination of energetics of scrunching and anti-scrunching

Lifetimes of unwound states (t_unwoun_d) were extracted from single-molecule traces as described (Revyakin et al., 2003, 2004, 2005, 2006) (Fig. 5A). Mean lifetimes of unwound states (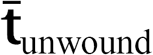) were extracted from histograms of t_unwoun_d, which exhibited single-exponential decay distributions, with means equal to standard deviations (Fig. 5B).

For experiments in absence of primer (Revyakin, 2004; Fig. S6):

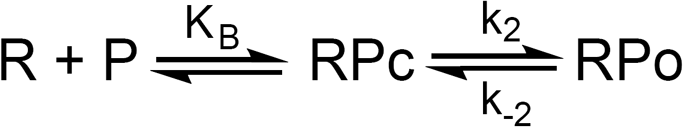

where R, P, RPc, and RPo denote RNAP holoenzyme, promoter, RNAP-promoter closed complex, and RNAP-promoter open complex, and

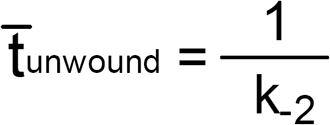

For experiments in presence of primer GGA, which programs TSS selection at modal position and therefore does not require scrunching or anti-scrunching for TSS selection (Fig. S6):

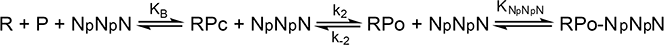

where NpNpN denotes primer; and

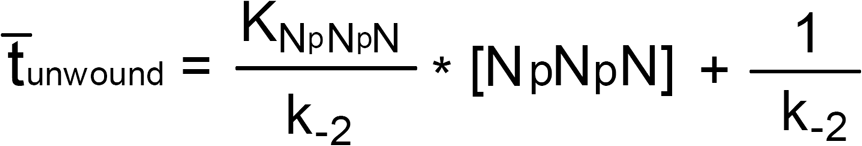

For experiments in presence of primer UGG, or GAA, or AAU, which program TSS selection at positions different form modal TSS, and therefore require scrunching or anti-scrunching for TSS selection (Figs. 4–6 and S6):

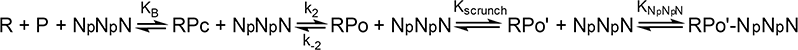

where RPo′ denotes a scrunched or anti-scrunched RPo; and

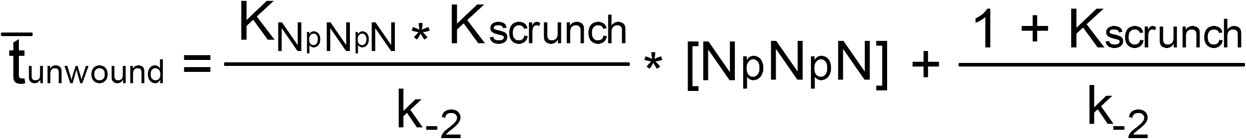

K_scrunch_, K_NpNpN_, ΔG_NpNpN_ and ΔG_scrunch_ were obtained by fitting slopes and y-intercepts of linear-regression fits of μlots of 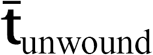 vs. primer concentration (Figs. 6D and S6) to the equation of Fig. 6C, stipulating K_scrunch_= 1 and ΔG_scrunch_= 0 for primer GGA, which programs TSS selection at modal position and therefore does not require scrunching or anti-scrunching for TSS selection.

## QUANTITATION AND STATISTICAL ANALYSIS

Data in Fig. 2 are means±SEM of at least 70 technical replicates of each of three biological replicates (three single DNA molecules) for positively supercoiled DNA and at least 50 technical replicates of each of three biological replicates (three single DNA molecules) for negatively supercoiled DNA.

Data in Fig. S4A-B are means±SEM of at least 100 technical replicates for a single DNA molecule (positively supercoiled DNA) or at least 70 technical replicates for a single DNA molecule (negatively supercoiled DNA).

Data in Fig. S4C are means±SEM of randomly selected subsets of n = 30. Similar results were obtained for ten different randomly selected subsets of n = 30.

Data in Figs. 4B-C and S5 are means±SEM of at least 40 technical replicates for each of seven biological replicates (seven single DNA molecules) for positively supercoiled DNA and at least 50 technical replicates for each of two biological replicates (two single DNA molecules) for negatively supercoiled DNA.

Data in Figs. 5–6 and S6 are means±SEM of at least 150 technical replicates for one single DNA molecule for each of the four primers.

## SUPPLEMENTARY FIG. LEGENDS

**Fig. S1.**
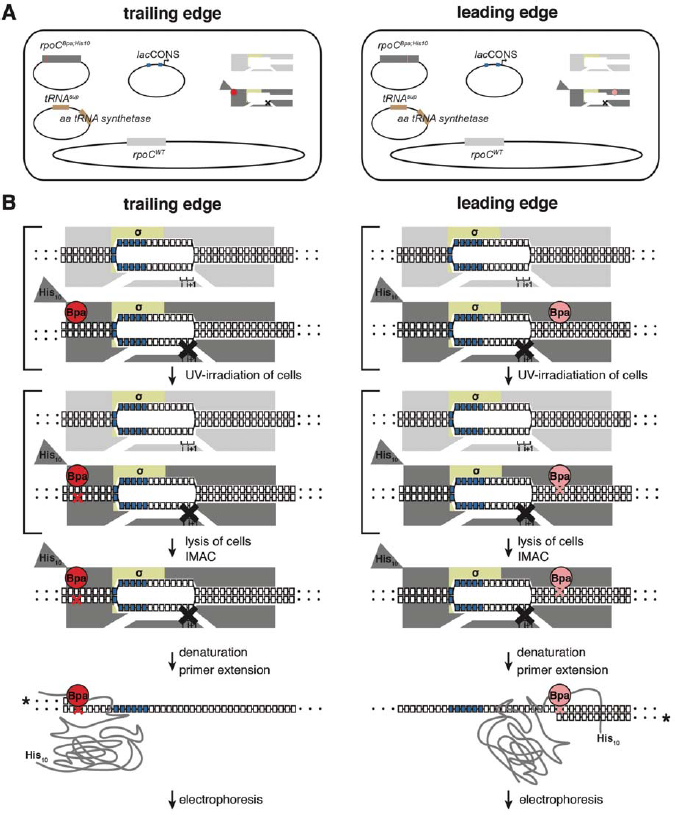
Unnatural-amino-acid mutagenesis and protein-DNA photocrosslinking *in vivo.* Left column, procedure for mapping RNAP trailing edge relative to DNA *in vivo.* Right column, procedure for mapping RNAP leading edge relative to DNA *in vivo.* (**A**) Three-plasmid merodiploid system for co-production, in *E. coli* cells, of Bpa-labeled, decahistidine-tagged, mutationally inactivated, RNAP-derivative (gray, with red or pink circle indicating Bpa, triangle indicating decahistidine tag, and black X indicating mutational inactivation), in presence of unlabeled, untagged, wild-type RNAP (light gray). First plasmid carries gene for RNAP subunit with a nonsense (TAG) codon at the site for incorporation of Bpa (residue β′T48 for trailing edge, residue β′R1148 for leading edge); second plasmid carries genes for engineered Bpa-specific nonsense-supressor tRNA and Bpa-specific amino-acyl-tRNA synthetase; third plasmid carries promoter of interest; and chromosome carries wild-type RNAP subunit genes. (**B**) Procedure for protein-DNA photocrosslinking, entailing UV-irradiation of cells, lysis of cells, immobilized-metal-ion affinity chromatography (IMAC), denaturation, primer extension with ^32^P-5’-end-labeled primer (asterisk denotes label), electrophoresis, and storage-phosphor imaging. Brackets represent steps performed in cells.

**Fig. S2.**
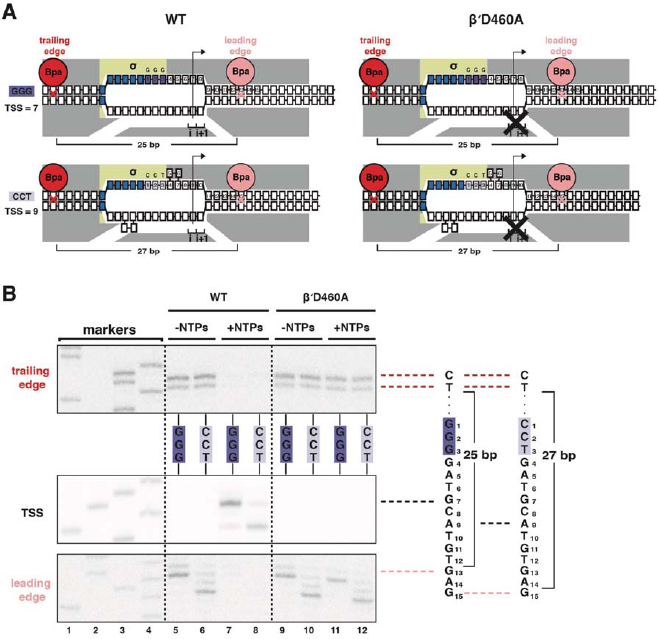
Mutationally inactivated RNAP derivative traps RPo in presence of NTPs, enabling protein-DNA photocrosslinking of RPo in presence of NTPs. (**A**) RNAP leading-edge and trailing-edge positions for wild-type RNAP (WT; left) and mutationally inactivated RNAP (β′D460A; right) at promoters having GGG discriminator (TSS 7 bp downstream of − 10 element; top) and CCT discriminator (TSS 9 bp downstream of −10 element; bottom). Black X, mutational inactivation due to β′D460A substitution. Other colors as in Fig. 1A. (**B**) RNAP trailing-edge crosslinking (top), TSS (middle), and RNAP leading-edge crosslinking (bottom) for wild-type RNAP (WT; left) and mutationally inactivated RNAP (β′D460A; right) at promoters having GGG discriminator and CCT discriminator, in absence and presence of NTPs. Horizontal dashed lines relate bands on gel (left) to nucleotide sequences (right).

**Fig. S3.**
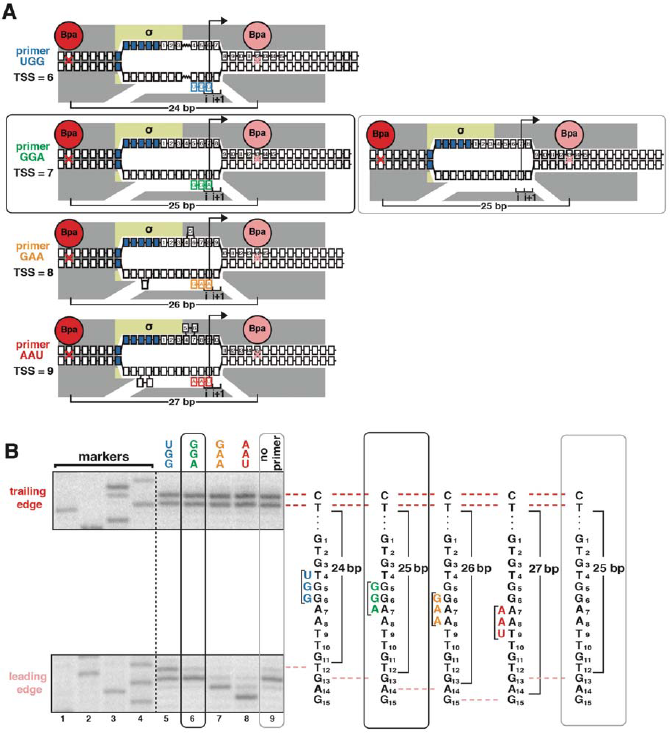
Protein-DNA photocrosslinking in primer-programmed TSS selection: primer GGA yields same pattern of RNAP leading-edge and trailing-edge crosslinking as in absence of primer. (**A**) RNAP leading-edge and trailing-edge positions and RNAP-dependent DNA unwinding with primers programming TSS selection at positions 6, 7, 8, and 9 bp downstream of −10 element (left panel), and with no primer (right panel). Black rectangle with rounded corners highlights case of primer GGA, which programs TSS selection at position 7. Gray rectangle with rounded corners highlights case of no primer, which also yields TSS selection at position 7. (**B**) Use of protein-DNA photocrosslinking to define RNAP leading-edge and RNAP trailing-edge positions. RNAP trailing-edge crosslinking (top), and RNAP leading-edge crosslinking (bottom) with primers UGG, GGA, GAA, and AAU (lanes 5-8), and with no primer (lane 9). Horizontal dashed lines relate bands on gel (left) to nucleotide sequences (right).

**Fig. S4.**
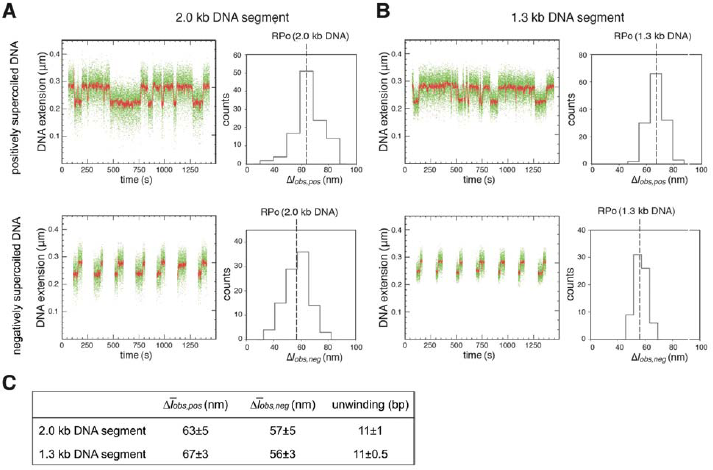
Single-molecule DNA-nanomanipulation: shorter DNA fragment enables detection of RNAP-dependent DNA unwinding with single-bp resolution. (**A-B**) Single-molecule time traces and transition-amplitude histograms with 2.0 kb DNA fragment [fragment length in previous work (Revyakin et al., 2005, 2006) and in Fig. 2] (**A**), and with 1.3 kb DNA fragment (fragment length in Figs. 4 and S5) (**B**). Data for positively supercoiled DNA are at top; data for negatively supercoiled DNA are at bottom. Green points, raw data (30 frames/s); red points, averaged data (1-s window). (**C**) Mean transition amplitudes on positively supercoiled DNA 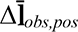, mean transition amplitudes on negatively supercoiled DNA(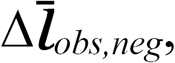), and DNA unwinding (means±SEM). Use of 1.3 kb DNA segment provides SEM = 0.5 bp for data subsets of n = 30, sufficient for single-bp resolution.

**Fig. S5.**
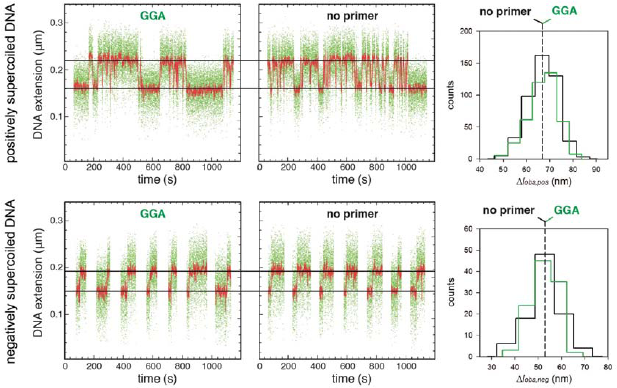
Single-molecule DNA-nanomanipulation: analysis of primer-programmed TSS selection with primer GGA. Single-molecule time traces and transition-amplitude histograms with GGA, which programs TSS selection 7 bp downstream of the −10 element, and and with no primer, which also yields TSS selection 7 bp downstream of the −10 element (data for positively supercoiled DNA at top; data for negatively supercoiled DNA below). Primer GGA results in same RNAP-dependent DNA unwinding as in absence of primer.

**Fig. S6.**
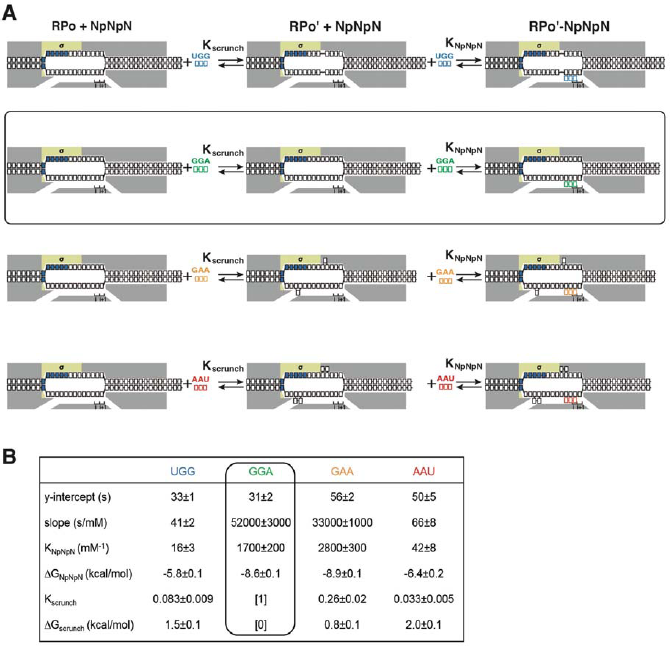
Single-molecule DNA-nanomanipulation: calculation of K_scrunch_ and ΔG_scrunch_ in primer-programmed TSS selection with primers UGG, GGA, GAA, and AAU. (**A**) Coupled equilibria for promoter scrunching or anti-scrunching (K_scrunch_) and primer binding (K_NpNpN_) with primers UGG, GGA, GAA, and AAU. Rectangle with rounded corners highlights case of primer GGA, which programs TSS selection corresponding to modal TSS in absence of primer (7 bp downstream of −10 element). Colors as in Fig. 3A.# (**B**) Intercepts and slopes of linear-regression μlots of primer-concentration dependence of 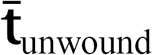 (lines 1-2; Fig. 6D), and K_NpNpN_, ΔG_NpNpN_, K_scrunch_, and ΔG_scrunch_ calculated from intercepts and slopes (lines 3-6; calculated using equation of Fig. 6C and 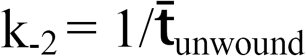 in absence of primer) (Revyakin et al., 2004).

**Table S1.**
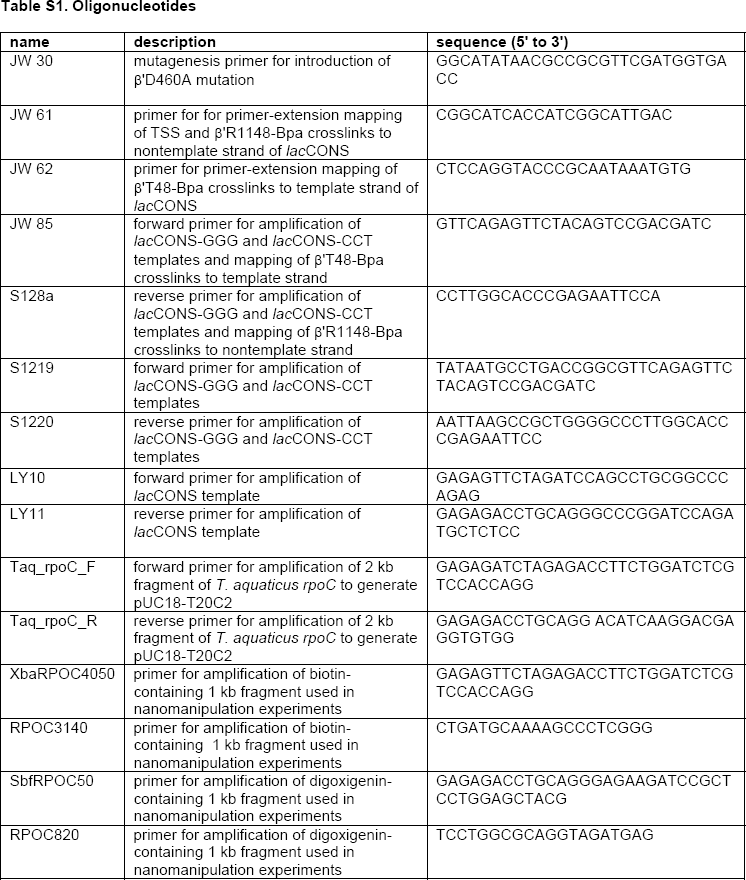
Oligonucleotides.

## References

Artsimovitch, I., Svetlov, V., Murakami, K.S., and Landick, R. (2003). Co-overexpression of *Escherichia coli* RNA polymerase subunits allows isolation and analysis of mutant enzymes lacking lineage-specific sequence insertions. J. Biol. Chem. 278, 12344–12355.

Bustamante, C., Marko, J., Siggia, E., and Smith, S. (1994). Entropic elasticity of lambda-phage DNA. Science 265, 1599–1600.

Chin, J.W., Martin, A.B., King, D.S., Wang, L., and Schultz, P.G. (2002). Addition of a photocrosslinking amino acid to the genetic code of *Escherichia coli*. Proc. Natl. Acad. Sci. USA. 99, 11020–11024.

Fazal, F., Meng, C., Murakami, K., Kornberg, R., and Block, S. (2015). Real-time observation of the initiation of RNA polymerase II transcription. Nature 525, 274–277.

Franklin, R., and Gosling, R. (1953). Molecular configuration in sodium thymonucleate. Nature 171, 740–741.

Feklistov, A., Barinova, N., Sevostyanova, Heyduk, E., Bass, I., Vvedenskaya, I., Kuznedelov, K., Merkiene, E., Stavroskaya, E., Klimasauskas, S., Nikiforov, V., Heyduk, T., Severinov, K., and Kulbachinsky, A. (2006). A basal promoter element recognized by free RNA polymerase sigma subunit determines promoter recognition by RNA polymerase holoenzyme. Mol. Cell 23, 97–107.

Fruscoloni, P., Zamboni, M., Panetta, G., De Paolis, A., and Tocchini-Valentinl, G. (1995). Mutational analysis of the transcription start site of the yeast tRNA Leu3gene. Nucleic Acids Research 23, 2914–2918.

Giardina, C., and Lis, J. (1993). DNA melting on yeast RNA polymerase II promoters. Science 261, 759–762.

Goldman, S., Sharp, J., Vvedenskaya, I., Livny, J., Dove, S., and Nickels, B. (2011). NanoRNAs prime transcription initiation *in vivo*. Mol. Cell 42, 817–825.

Haugen, S., Berkmen, M., Ross, W., Gaal, T., Ward, C., and Gourse, R. (2006). rRNA promoter regulation by nonoptimal binding of sigma region 1.2: an additional recognition element for RNA polymerase. Cell 125, 1069–1082.

Jeong, W., and Kang, C. (1994). Start site selection at lacUV5 promoter affected by the sequence context around the initiation sites. Nucleic Acids Res. 22, 4667–4672.

Kuehner, J., and Brow, D. (2006). Quantitative analysis of *in vivo* initiator selection by yeast RNA polymerase II supports a scanning model. J. Biol. Chem. 281, 14119–14128.

Learned, R., and Tjian, R. (1982). *In vitro* transcription of human ribosomal RNA genes by RNA polymerase I. J. Mol. Appl. Genet. 1, 575–584.

Lewis, D.E., and Adhya, S. (2004). Axiom of determining transcription start points by RNA polymerase in Escherichia coli. Mol. Micro. 54, 692–701.

Lionnet, T., Allemand, J., Revyakin, A., Strick, T., Saleh, O., Bensimon, D., and Croquette, V. (2012). Magnetic trap construction. Cold Spring Harbor Protocols 2012, 133–138.

Liu, J., and Turnbough, C.L. (1994). Effects of transcriptional start site sequence and position on nucleotide-sensitive selection of alternative start sites at the *pyrC* promoter in *Escherichia coli*. J. Bacteriol. 176, 2938–2945.

Marr, M., and Roberts, J. (1997). Promoter recognition as measured by binding of polymerase to nontemplate strand oligonucleotide. Science 276, 1258–1260.

Mukhopadhyay, J., Kapanidis, A., Mekler, V., Kortkhonjia, E., Ebright, Y., and Ebright, R. (2001). Translocation of sigma(70) with RNA polymerase during transcription: fluorescence resonance energy transfer assay for movement relative to DNA. Cell 106, 453–463.

Reiter, W., Hüdepohl, U., and Zillig, W. (1990). Mutational analysis of an archaebacterial promoter: essential role of a TATA box for transcription efficiency and start-site selection *in vitro*. Proc. Natl. Acad. Sci. U.S.A. 87, 9509–9513.

Revyakin, A., Allemand, J., Croquette, V., Ebright, R., and Strick, T. (2003). Single-molecule DNA nanomanipulation: detection of promoter-unwinding events by RNA polymerase. Meths. Enzymol. 370, 577–598.

Revyakin, A., Ebright, R., and Strick, T. (2004). Promoter unwinding and promoter clearance by RNA polymerase: detection by single-molecule DNA nanomanipulation. Proc. Natl. Acad. Sci. U.S.A. 101, 4776–4780.

Revyakin, A., Ebright, R., and Strick, T. (2005). Single-molecule DNA nanomanipulation: Improved resolution through use of shorter DNA fragments. Nature Methods 2, 127–138.

Revyakin, A., Liu, C., Ebright, R., and Strick, T. (2006). Abortive initiation and productive initiation by RNA Polymerase involve DNA scrunching. Science 314, 1139–1143.

Robb, N., Cordes, T., Hwang, L., Gryte, K., Duchi, D., Craggs, T., Santoso, Y., Weiss, S., Ebright, R., and Kapanidis, A. (2013). The transcription bubble of the RNA Polymerase–Promoter Open Complex Exhibits Conformational Heterogeneity and Millisecond-Scale Dynamics: Implications for Transcription Start-Site Selection. J. Mol. Biol. 425, 875–885.

Sainsbury, S., Bernecky, C., and Cramer, P. (2015). Structural basis of transcription initiation by RNA polymerase II. Nat. Rev. Mol. Cell Biol. 16, 129–143.

Sambrook, J., and Russell, D. (2001). Molecular Cloning: A Laboratory Manual (Cold Spring Harbor Laboratory, Cold Spring Harbor, NY).

Samuels, M., Fire, A., and Sharp, P. (1984). Dinucleotide priming of transcription mediated by RNA polymerase II. J. Biol. Chem. 259, 2517–2525.

Svetlov, V., and Artsimovitch, I. (2015). Purification of bacterial RNA polymerase: tools and protocols. Methods Mol Biol. 1276, 13–29.

Thomm, M., and Wich, G. (1988). An archaebacterial promoter element for stable RNA genes with homology to the TATA box of higher eukaryotes. Nucleic Acids Research 16, 151–163.

Vvedenskaya, I., Zhang, Y., Goldman, S., Valenti, A., Visone, V., Taylor, D., Ebright, R., and Nickels, B. (2015). Massively systematic transcript end readout, "MASTER": transcription start site selection, transcriptional slippage, and transcript yields. Mol. Cell 60, 953–965.

Walker, K., and Osuna, R. (2002). Factors affecting start site selection at the *Escherichia coli fis* promoter. J. Bacteriol. 184, 4783–4791.

Wang, J. C. (1979). Helical repeat of DNA in solution. Proc. Natl. Acad. Sci. U.S.A. 76, 200–203.

Winkelman, J., Winkelman, B., Boyce, J., Maloney, M., Chen, A., Ross, W., and Gourse, R. (2015). Crosslink mapping at amino acid-base resolution reveals the path of scrunched DNA in initial transcribing complexes. Mol. Cell 59, 768–780.

Winkelman, J., Vvedenskaya, I., Zhang, Y., Zhang, Y., Bird, J., Taylor, D., Gourse, R., Ebright, R., and Nickels, B. (2016a). Multiplexed protein-DNA cross-linking: Scrunching in transcription start site selection. Science 351, 1090–1093.

Winkelman, J., Chandrangsu, P., Ross, W., and Gourse, R. (2016b). Open complex scrunching before nucleotide addition accounts for the unusual transcription start site of *E. coli* ribosomal RNA promoters. Proc. Natl. Acad. Sci. U.S.A. 113, e1787–e1795.

Zaychikov, E., Martin, E., Denissova, L., Kozlov, M., Markovtsov, V., Kashlev, M., Heumann, H., Nikiforov, V., Goldfarb, A., and Mustaev, A. (1996). Mapping of catalytic residues in the RNA polymerase active center. Science 273, 107–109.

Zecherle, G., Whelen, S., and Hall, B. (1996). Purines are required at the 5′ ends of newly initiated RNAs for optimal RNA polymerase III gene expression. Mol. Cell. Biol. 16, 5801–5810.

